# Circular mRNA against CleanCap linear mRNA vectors: comprehensive comparison, expression, active and passive immunization

**DOI:** 10.1101/2025.10.13.682008

**Authors:** Vladimir M. Vakhtinskii, Irina L. Tutykhina, Alina S. Dzharullaeva, Daria M. Grousova, Ilya D. Zorkov, Anna A. Ilyukhina, Dmitrii A. Reshetnikov, Valentin V. Azizyan, Artem A. Derkaev, Evgeniia N. Bykonia, Evgeny V. Usachev, Denis Kleymenov, Vladimir A. Gushchin, Inna V. Dolzhikova, Dmitry V. Shcheblyakov, Maxim M. Shmarov, Denis Yu. Logunov, Alexander L. Gintsburg

## Abstract

The mRNA platform has revolutionised vaccine technology by providing a universal, fast and easily scalable production process. There are two main types of mRNA – linear (cap-dependent) and circular (cap-independent), each with its own advantages. Although both vector types are continuously being improved, there has been no comprehensive comparative analysis of the most efficient existing vectors of each type. This paper provides a comparison of expression efficiency, protective and therapeutic activities of circular and linear mRNA vectors. We examined different combinations of linear vectors: containing either N1-methylpseudouridines or uridine and capped with either ARCA (m^7^G(5′)ppp(5′)G) or CleanCap (m**^7^**G(5′)ppp(5′)m^2^G). Circular vectors contained both commonly used IRES of coxsackievirus B3 and new a IRES of human rhinoviruses B6.

Preliminary luciferase assay showed that modified linear vectors exhibited significantly higher expression levels both *in vitro* and *in vivo*. A similar but less dramatic difference was observed in expression of the target protein. At the same time, immunogenicity and protective activity of linear and circular vectors were equal.

## Introduction

mRNA-based vaccines have already demonstrated a high level of safety and protective activity during SARS-CoV-2 pandemic [1].

In comparison to traditional vaccines, the mRNA platform offers a number of benefits. Among these are a standardized technological process, independent of the target pathogen, which facilitates large-scale production and simplifies regulatory approval [2].

There are two main types of mRNA vectors: linear (cap-dependant) and circular (cap independent). Both have their advantages and disadvantages. Linear vectors are better studied and more used in vaccine production. One of the best examples is a widely used vaccine against SARS-CoV-2 which is based on cap-dependent mRNA technology [1,3,4].

Circular mRNA vectors are relatively new, nevertheless, in recent years a number of vaccines based on this technology have showed their immunogenicity against various viral pathogens, including the Zika virus, Influenza viruses, Rabies-virus, and Monkeypox virus. Finally, circular mRNA was used as a platform for cancer vaccine [5,6,7,8,9].

The efficacy of circular mRNA is mainly determined by the Internal Ribosome Entering Site (IRES). There is a great diversity of known IRESes, but most of them are poorly studied and not used routinely. A set of frequently used IRESes, originates from human rhinoviruses A and B (HRV-A1. HRV-B3), coxsackievirus B3 (CVB3), encephalomyocarditis virus (EMCV), cricket paralysis virus (CrPV), poliovirus 1 (PV1) and hepatitis C virus (HCV). Among these the CVB3 IRES is a major one [10].

Expression levels of linear mRNA-vectors are based on the cap type and chosen 5′ and 3′ untranslated regions (UTRs). Cap is a structure at 5’ end of mRNA of eukaryotes. There are two main types of natural cap structures cap0 – 7-methylguanosine, linked by 5’-5’-triphosphate bridge (m7GpppNp) and cap1 – which contains additional methyl group in the first nucleotide (m7GpppNm2’Op).

There is an *in vitro* capping technology, providing single-step capping with cap0 type and an optional second stage for the enzymatic formation of cap1. Nevertheless, this approach is rather expensive. Another option is a use of cap analogues, such as Anti-Reverse Cap Analogue (ARCA). This technology offers a single-step co-transcriptional capping; however, its efficiency is about 80%. By now, the most effective and widely used capping reagent is m7G(5′)ppp(5′)(2’OMeA)pG also known as CleanCap AG. As an example, it was used for production of BNT162b2 vaccine against SARS-CoV2.

Important role belongs to UTRs, certain researches investigate natural UTRs, others screen the libraries of artificial UTRs. The most widely used and effective UTRs to the date are UTRs of alfa-globin, that were applied in BNT162b2 SARS-Cov2 vaccine [11,12,13,14].

A number of studies are devoted to the comparison of luciferase activity of circular and linear vectors, but fewer studies have provided immunogenicity and protective activity data. Among them are the work of Wan, J et all, and the work of Qu L. et.al. Wan, J et. al compared the immunogenicity of linear and circular vectors on the model of surface glycoprotein G of Rabies virus. And Qu L. et.al. studied used as a model antigen RBD-domain of S-glycoprotein of SARS-Cov2 [15,16].

Interesting, that both Wan, J et all and Qu L. et.al. used linear vectors, capped with first-generation cap analogues (m7G (5′) ppp (5′) G), while the approved linear mRNA-vaccines use more effective third-generation CleanCap reagent (m7G (5′) ppp (5′) m2G) [17].

That is why the aim of our study was to compare immunogenicity and protective activity of different linear mRNA capped with various cap types against circular mRNA vector containing different IRESes. As an etalon of UTRs of linear mRNA vectors we chose the UTRs of alfa-globin, as they are considered among the most efficient and safe.

Recent research claims that the IRESes of some Rhinoviruses are more effective than CVB3. Therefore, in our work, in addition to “popular” IRES CVB3 we also used the yet-not studied IRES of Rhinovirus B6 (HRV B6) [18].

## Results

### Linear and circular mRNA-vectors

The synthesis scheme of linear and circular vectors is shown in Figure 1. All the DNA templates contain T7 promoter, which is recognized by T7 mRNA-polymerase. Two linear templates have 5′and 3′ untranslated regions (5′UTR and 3′UTR) from alfa-globin, the gene of interest, and poly-(A) sequence, consisting of 30 and 70 residuals separated by a 10-nucleotide “island”. The difference between the two linear vectors lies in the two codons after the T7 promoter. These codons define the capping system (AG was used for CleanCap AG reagent, while GG was used for ARCA-capping). Both linear vectors were made with uridine or with its substitution with N1-methylpseudouridine. Circular vectors have special permuted intron-exons (PIE) required for mRNA cyclization at the 5′ and 3′ ends of the mRNA IRES and the gene of interest. The distinction between them is in the IRES used: CVB3 or HRVB6.

**Fig. 1.**
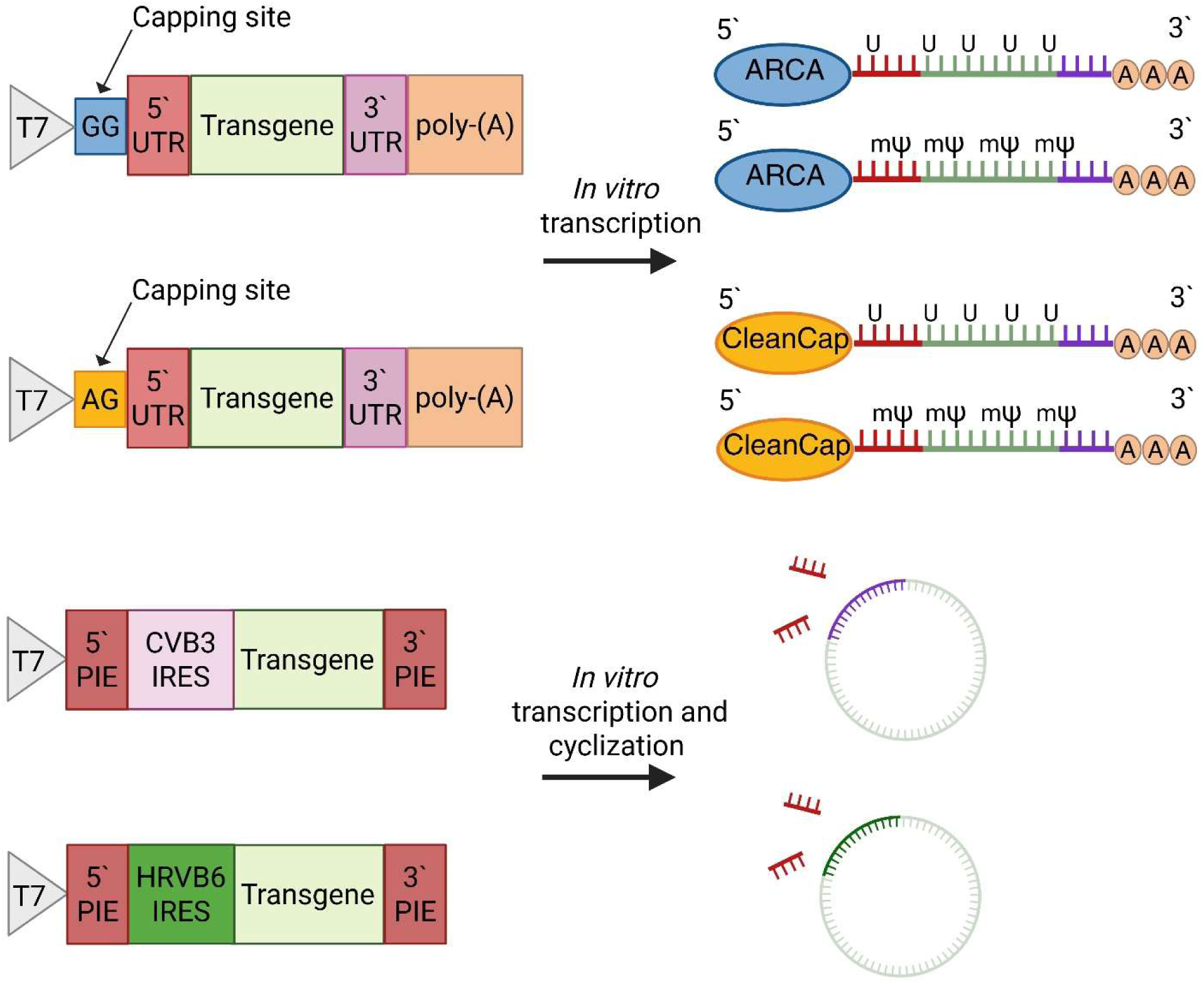
schematic process of mRNA-vectors production. Linear vectors contained either uridine or N1-methylpseudouridine and were capped with ARCA reagent or CleanCap AG reagents. Circular vectors include either CVB3 or HRVB6 IRES and did not contain N1-methylpseudouridine.

Linear vectors were produced using a single-step procedure of mRNA-capping, which occurs co-transcriptionally. In contrast, circular vectors required a second step of cyclization followed by the treatment with RNase R to remove introns and linear precursors.

### *In vitro* luciferase activity

The first step of our experiment was to compare the *in vitro* expression of different constructs using a firefly luciferase assay. HEK-293 cells were transfected with different mRNA-vectors. The 120-hour-long kinetic curves of luciferase activity are presented in Figure 2. Values of luciferase activity at selected time-points are presented in Table 1.

**Figure 2.**
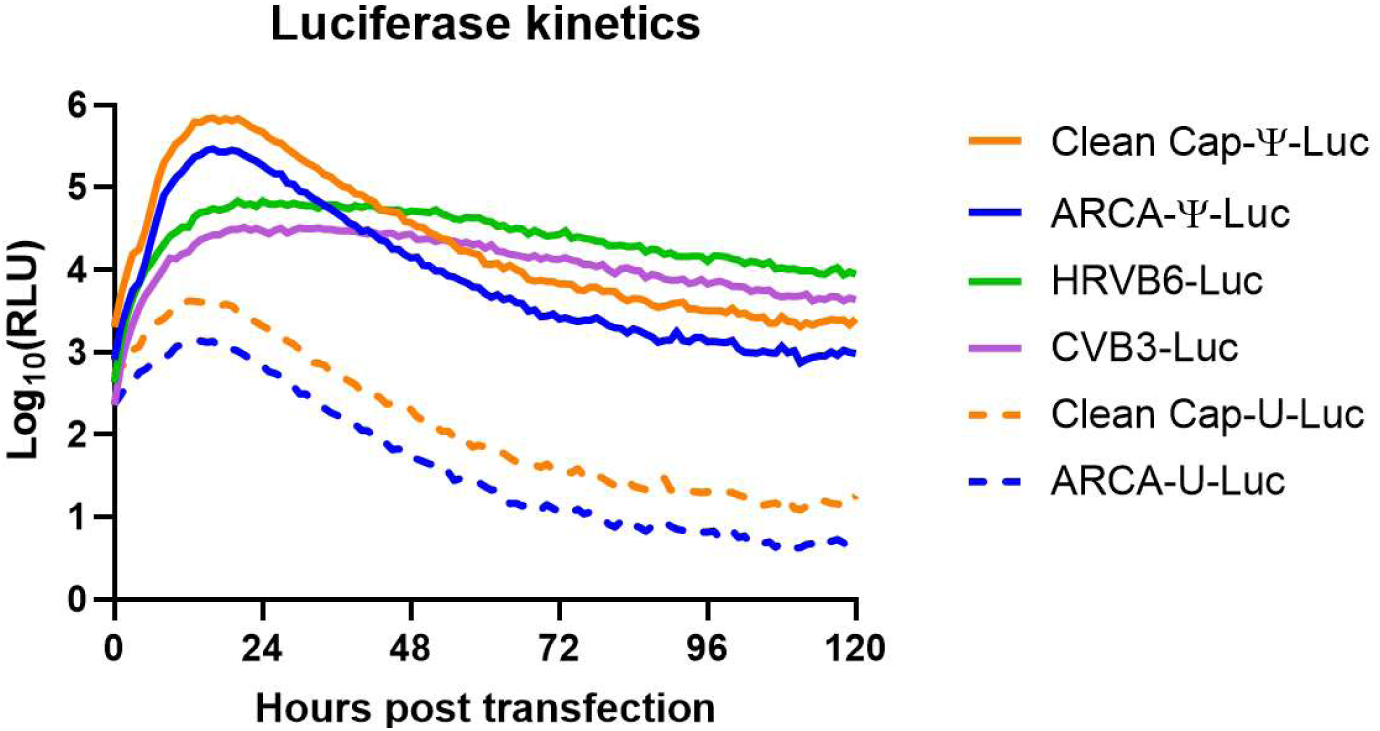
*In vitro* kinetics of luciferase activity at HEK-293 cell line. (The experiment was performed three times at different cell passages; for each experiment all the mRNA-vectors were freshly prepared.

**Table 1.** *In vitro* luciferase activity at selected time-points (mean with CI 95%)

| **mRNA-**<br>**vector** | **10 hours** | **24 hours** | **48 hours** | **72 hours** | **96 hours** | **120 hours** |
| --- | --- | --- | --- | --- | --- | --- |
| CleanCap Ψ | 340 144.3<br>±29 534.5 | 472 710.2<br>±17 678.9 | 37 323.7<br>±3 602.7 | 6 732.3<br>±251.8 | 3 193.7<br>±326.2 | 2 569.3<br>±157.6 |
| ARCA-Ψ | 130 494.3<br>±10 832.9 | 185 766.8<br>±16 258.3 | 138 76.1<br>±1 698.3 | 2 517.6<br>±258.9 | 1 345.9<br>±63.8 | 964.8<br>±67.7 |
| HRVB6 | 28 538.7<br>±2 605.1 | 68 002.1<br>±5 582.4 | 51 310.4<br>±4 404.4 | 26 724.8<br>±2 696.7 | 12 672.5<br>±953.7 | 8 849.6<br>±872.7 |
| CVB3 | 13 530.4<br>±1 245.8 | 29 550.2<br>±2 899.5 | 27 071.6<br>±1 804.4 | 13 430.3<br>±1 405.7 | 6 752.5<br>±680.3 | 4 328.6<br>±490.9 |
| CleanCap-U | 3 735.7<br>±401.7 | 2 075.3<br>±244.9 | 202.9<br>±13.5 | 35.6<br>±3.7 | 20<br>±1.7 | 18.3<br>±1 |
| ARCA-U | 1 166.8<br>±89.1 | 694.2<br>±39 | 62.0<br>±5.2 | 11.7<br>±1.1 | 6.5<br>±0.6 | 5.3<br>±0.6 |

As shown, the peak of luciferase activity for linear vectors occurs between 16 and 24 hours, while for “circular” vectors the maximum is observed at 24 hours.

Linear vectors containing N1-methylpseudouridine demonstrated significantly higher luciferase activity compared to circular vectors. At the same time linear non-modified vectors exhibit low luciferase activity and a more dramatic decline over the time. By 24 hours the activity of non-modified linear vectors had decreased more than that of modified vectors.

The highest luciferase activity was observed for the linear vector capped with CleanCap reagent and modified with N1-methylpseudouridine at the 24- and 10-hour time-points: 472 710.2 ±17 678.9 and 340 144.3 ± 29 534.5 respectively. The second highest activity was observed in a linear modified vector but capped with ARCA reagent, at the same time points: 185 766.8 ± 16 258.3 and 130 494.3 ± 10 832.9 respectively.

The circular vector with HRVB6 IRES ranked third with luciferase activity 68 002.1 ± 5 582.4 at 24-hour time point. The circular vector with CVB3 IRES was fourth with luciferase activity 29 550.2 ± 2 899.5 at 24-hour time point. Linear non-modified vectors showed poor results, not exceeding 4000.

Based on these results of the *in vivo* luciferase activity, we excluded both linear mRNA with uridine from further *in vivo* experiments as they was the least effective in human considerations.

### *In vivo* luciferase activity

BALB/c female mice were injected intramuscularly with lipid nanoparticles (LNPs), containing different luciferase-coding vectors. The dose of mRNA was 10 µg per mouse. At various time points, luciferase activity was measured using an IVIS Lumina II (Life sciences). The visualised results are shown in Figure 3 and quantitative data is presented in Table 2.

**Figure 3.**
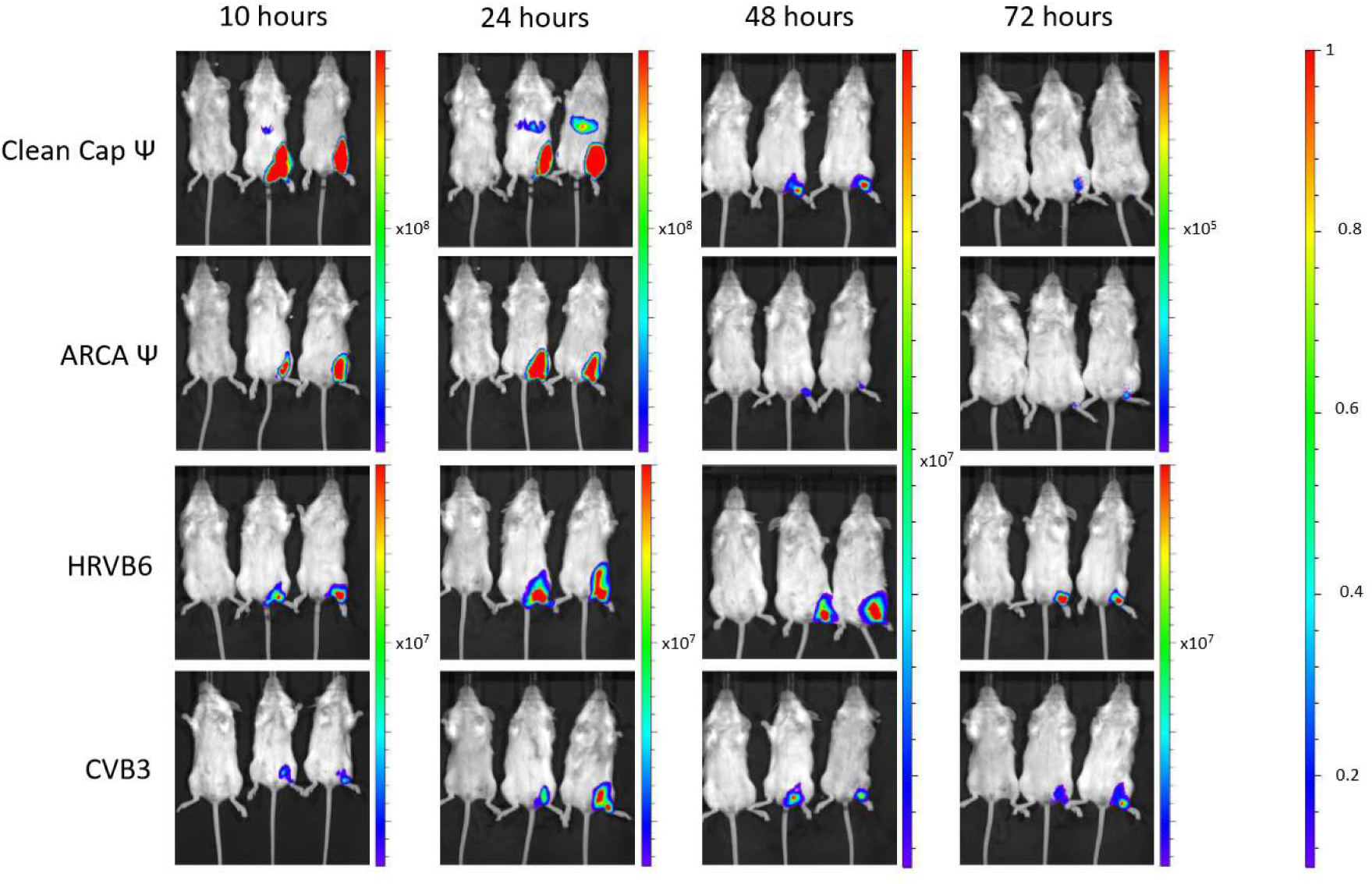
*In vivo* luciferase activity in BALB/c mice at various time-points.

**Table 2.** Total calculated *in vivo* luciferase activity, [p/s].

| Vector | mice | Total Flux [p/s] |  |  |  |
| --- | --- | --- | --- | --- | --- |
|  |  | 10 hours | 24 hours | 48 hours | 72 hours |
| CleanCap Ψ | 1 | 6.42x10<sup>9</sup> | 7.62x10<sup>9</sup> | 5.25 x10<sup>7</sup> | 2.50 x10<sup>5</sup> |
|  | 2 | 4.19 x 10<sup>9</sup> | 5.70X10<sup>9</sup> | 4.22 x10<sup>7</sup> | 1.31 x10<sup>5</sup> |
|  | mock | 4.73 x10<sup>7</sup> | 8.69 x10<sup>7</sup> | 1.95 x10<sup>5</sup> | 1.07 x10<sup>5</sup> |
| ARCA Ψ | 1 | 1.73 x10<sup>9</sup> | 3.27x10<sup>9</sup> | 8.10 x10<sup>6</sup> | 1.09 x10<sup>5</sup> |
|  | 2 | 2.96 x10<sup>9</sup> | 4.39 x10<sup>9</sup> | 5.80 x10<sup>6</sup> | 1.64 x10<sup>5</sup> |
|  | mock | 4.42 x10<sup>6</sup> | 1.07 x10<sup>7</sup> | 1.45 x10<sup>5</sup> | 9.93 x10<sup>4</sup> |
| HRVB6 | 1 | 6.57 x10<sup>7</sup> | 5.27 x10<sup>8</sup> | 1.47 x10<sup>8</sup> | 7.67 x10<sup>7</sup> |
|  | 2 | 1.10 x10<sup>8</sup> | 5.75 x10<sup>8</sup> | 2.28 x10<sup>8</sup> | 1.18 x10<sup>8</sup> |
|  | mock | 2.53 x10<sup>5</sup> | 8.25 x10<sup>5</sup> | 3.53 x10<sup>5</sup> | 2.69 x10<sup>5</sup> |
| CVB3 | 1 | 2.74 x10<sup>7</sup> | 4.25 x10<sup>7</sup> | 4.43 x10<sup>7</sup> | 1.74 x10<sup>7</sup> |
|  | 2 | 2.61 x10<sup>7</sup> | 1.52 x10<sup>8</sup> | 2.34 x10<sup>7</sup> | 4.09 x10<sup>7</sup> |
|  | mock | 2.00 x10<sup>5</sup> | 2.79 x10<sup>5</sup> | 2.57 x10<sup>5</sup> | 1.93 x10<sup>5</sup> |

*In vivo* luciferase activity results were consistent with the *in vitro* findings. Mice treated with linear mRNA vector reached near-maximum values at 10 and 24 hours, followed by a dramatic decrease in luciferase activity over the next 24 hours. Circular vectors showed a slower decline in luciferase activity.

As *in vitro,* the highest total luciferase activity *in vivo* was observed for the linear vector capped with CleanCap reagent and modified with N1-methylpseudouridine at the 24-hour time-point. The second highest was the linear modified vector capped with ARCA reagent also at the 24-hour time-point. Circular vectors HRVB6 and CVB3 were ranked third and fourth respectively.

Our next step was to evaluate the expression levels of target protein on the model of the SARS-CoV-2 S glycoprotein. This gene was inserted into each of the vector types.

### Expression of S-glycoprotein of SARS-Cov2

HEK-293 cells in a 6-well plate with were transfected with different mRNA vectors. As an additional negative control, we used the most productive luciferase-coding vector (CleanCap-Ψ-Luc). Twenty hours post transfection, cells were washed and lysed. Cell lysate supernatants were used for ELISA against the S-glycoprotein of SARS-CoV2. The results are shown in Figure 4, and the mean values with 95% confidence intervals are presented in Table 3.

**Fig. 4.**
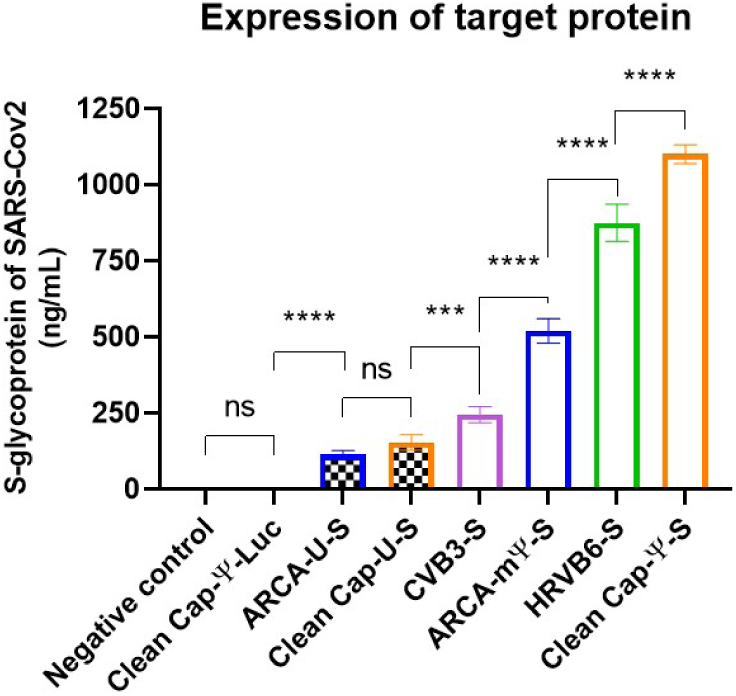
I*n vitro* expression of SARS-CoV2 S-glycoprotein in HEK-293 cells 20 hours post-transfection. Data is presented as a mean with 95% CI intervals. P-values were determined by the non-parametric ANOVA (Friedman’s test) with Dunn’s multiple comparison post-test ***p=0.0002; ****p<0.0001

**Table. 3.** Expression levels of SARS-CoV2 S-glycoprotein in HEK-293 cells (Mean and 95% CI)* ng/mL.

| Vector | ARCA-U-S | CleanCap<br>-U-S | CVB3-S | ARCA-Ψ-S | HRVB6-S | CleanCap-Ψ-S |
| --- | --- | --- | --- | --- | --- | --- |
| **Concentration**<br>**, ng/mL** | 114.3<br>±12.5 | 153.2<br>±26.45 | 244.7<br>±27.95 | 519.5<br>±42.1 | 875.0<br>±63.7 | 1100<br>±32 |

\* Zero means of luciferase-coding vector and negative control are not shown

As well as in the luciferase assay the highest expression was observed for the linear vector capped with CleanCap reagent and modified with N1-methylpseudouridine (1100 ± 32). Surprisingly the second-higher expression level belonged to the circular mRNA containing the HRVB6 IRES (875.0 ± 63.7). The linear modified vector capped with the ARCA reagent ranked third with 519.5 ±42.1 ng/ml. The circular vector with the CVB IRES showed 244.7 ± 27.95 ng/ml. Linear non-modified vectors exhibited poor expression: 153.2 ± 26.45 and 114.3 ± 12.5 for the CleanCap and for the ARCA respectively.

We selected the 20-hour time point for the *in vitro* protein-expression experiment with the luciferase assay. By the 20^th^ hour, the linear vectors remain on the plateau, while the circular vectors have just reached it, making this an optimal moment for comparing “maximum” expression levels.

Next, we investigated the immunogenicity of the generated vectors in mice. As during *in vivo* luciferase assay, we excluded linear vectors containing canonical nucleotides in human consideration, because of their instability.

### Immunogenicity study

The first step was a single-dose intramuscular immunization of ACE-2 mice with mRNA at a dose of 10 µg/mouse or PBS for the negative control. Each group contained 10 mice.

On day 21 post-immunization we measured an antibody titres against the RBD-domain of SARS-CoV-2 (Wuhan) in the serum of the vaccinated animals. Graphical results are shown in Figure 5 and the endpoint GMT is presented in Table 4.

**Figure 5.**
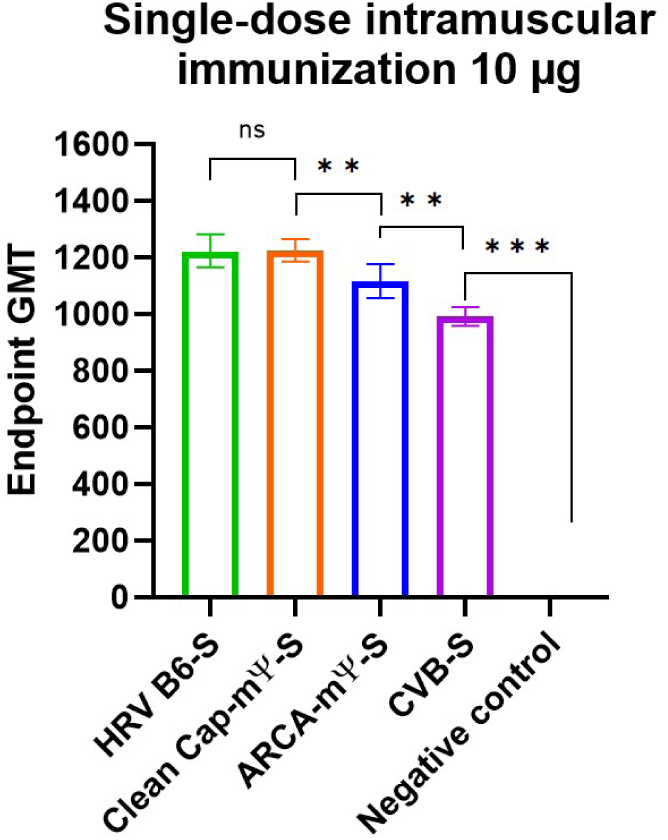
Endpoint GMT of anti-S-glycoprotein of SARS-CoV2 in serum of mice after a single-dose immunization at a dose of 10 µg per mouse. Data is presented as geometric mean with 95% CI intervals. P-values were determined by the Mann–Whitney test **p< 0.0078; ***p<0.0005

**Table. 4.** Endpoint GMT of anti-S-glycoprotein of SARS-CoV2 in serum of mice after a single-dose immunization at a dose of 10 µg per mouse. (Geometric mean and 95% CI)*

| Vector | HRVB6-S | CleanCap-Ψ-S | ARCA-Ψ-S | CVB3-S |
| --- | --- | --- | --- | --- |
| Endpoint GMT | 1222±83 | 1224±40 | 1113±60 | 990±48 |

\* Zero means of negative control is not shown

Results of a single-dose immunisation showed comparable endpoint GMTs for the linear vector capped with CleanCap reagent and modified with N1-methylpseudouridine and the circular vector with the HRVB6 IRES (1224 ± 40 and 1222 ± 83, respectively). The circular vector with the CVB3 IRES had the lowest endpoint GMT (990 ± 48), while the linear vector capped with the ARCA reagent and modified with N1-methylpseudouridine showed an endpoint GMT 1113 ± 60.

For subsequent work we excluded the two lowest-performing vectors, being a linear mRNA with ARCA and circular mRNA with CVB3 IRES. In the next experiment we used a classical prime-boost immunisation with the reduced doses (3 µg per mouse).

### Immunogenicity and protective activity of two “best” vectors

ACE-2 mice were divided into three groups with 30 mice in each. The immunization scheme remained the same, as previously described. On day 21 after the second immunization, 10 mice from each group were selected for blood collection and subsequent antibody titre measurement. The results are shown in Figure 6A.

**Figure 6.**
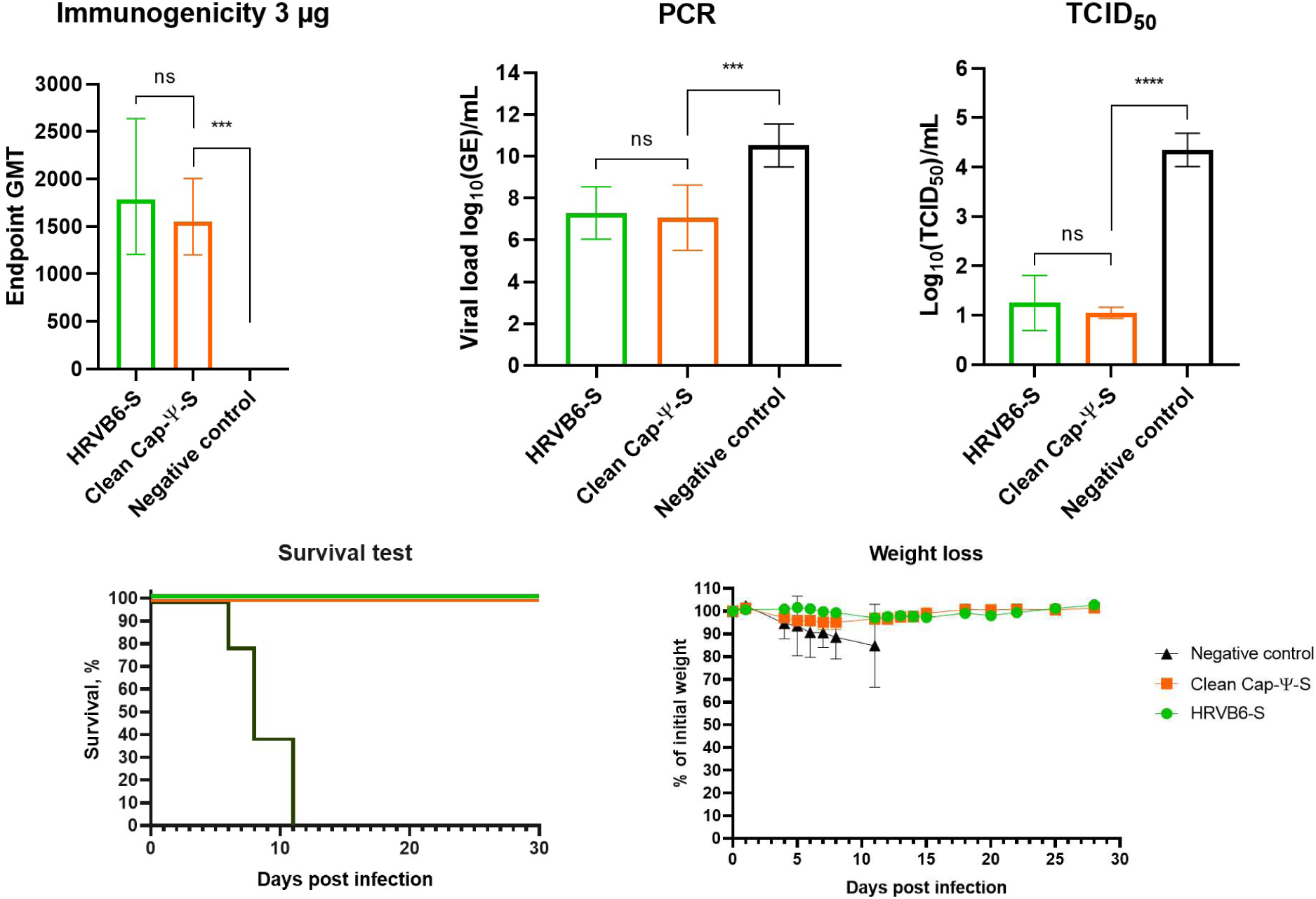
SARS-CoV2 experiment. (A) – Immunogenicity of circular and linear mRNA vectors after prime-boost vaccination at a dose 3 µg. (B) – Viral load measured by PCR-RT in lungs of immunized mice on day 3 post SARS-CoV2 challenge. (C) – Viral load measured by TCID_50_ titration in lungs of immunized mice on day 3 post SARS-CoV2 challenge. (D) – Survival after SARS-CoV2 challenge. (E) – Weight loss after SARS-CoV2 challenge.

The remaining mice were challenged with the SARS-CoV2 Wuhan strain at a dose 10^5^ TCID_50_. On the third day post-infection 10 mice from each group were sacrificed. Viral load in the lungs was measured using RT-PCR and TCID_50_ assay on Vero cells. The results are presented in Figure 6B and 6C. The remaining mice were monitored for four weeks with daily weight measurements (Figure 6D and 6E).

Both groups of immunized mice showed equivalent GMTs of anti-RBD neutralizing antibodies, significantly higher than the negative control (1783 ± 715 for the circular vector with HRVB6 IRES and 1552 ±402 for the linear CleanCap vector).

Viral load measured by RT-PCR also showed a significant decrease in viral load in lungs. Log_10_ of genome copies were 7.283 ± 1.251 for circular vector and 7.063 ±1.559 for the linear vector compared to 10.52 ±1.027 in the control group

TCID_50_ titration also indicated lower viral loads in vaccinated mice. Log_10_(TCID_50_) was 1.25±0.694 for the circular vector and 1.05±0.139 for the linear vector, versus 4.35±0.416 for the control group.

Survival test showed 100% protection from lethal outcome in both linear and circular vector-immunized groups, while all unvaccinated mice died by day11. The results of daily weigh monitoring indicated that all the mice lost weight during the first 11 days post-infection, after which vaccinated mice began re-gaining weight. Mice immunized with the linear vector experienced less weight loss, compared to those vaccinated with the circular vector. Nevertheless, by day 26 post-infection all vaccinated groups reached their initial weight.

### Application of circular mRNA-vector for passive immunization

To complete a multifaceted comparison, we evaluated the protective efficacy of circular mRNA-vectors expressing recombinant antibody against *C. botulinum* toxin A (BonT/A).

We generated a circular vector, containing IRES HRVB6 coding B11-Fc antibody. The linear vector capped with CleanCap reagent was described previously [15].

Mice were divided into three groups of 12 animals each and were treated with LNPs, containing either a linear or circular vector at a dose of 10 µg per mouse. As a negative control a luciferase-coding linear vector was used. At various timepoints the mice underwent blood collection and the subsequent recombinant antibody titre measurement in serum by ELISA. The results are shown in Figure 7.

**Figure 7.**
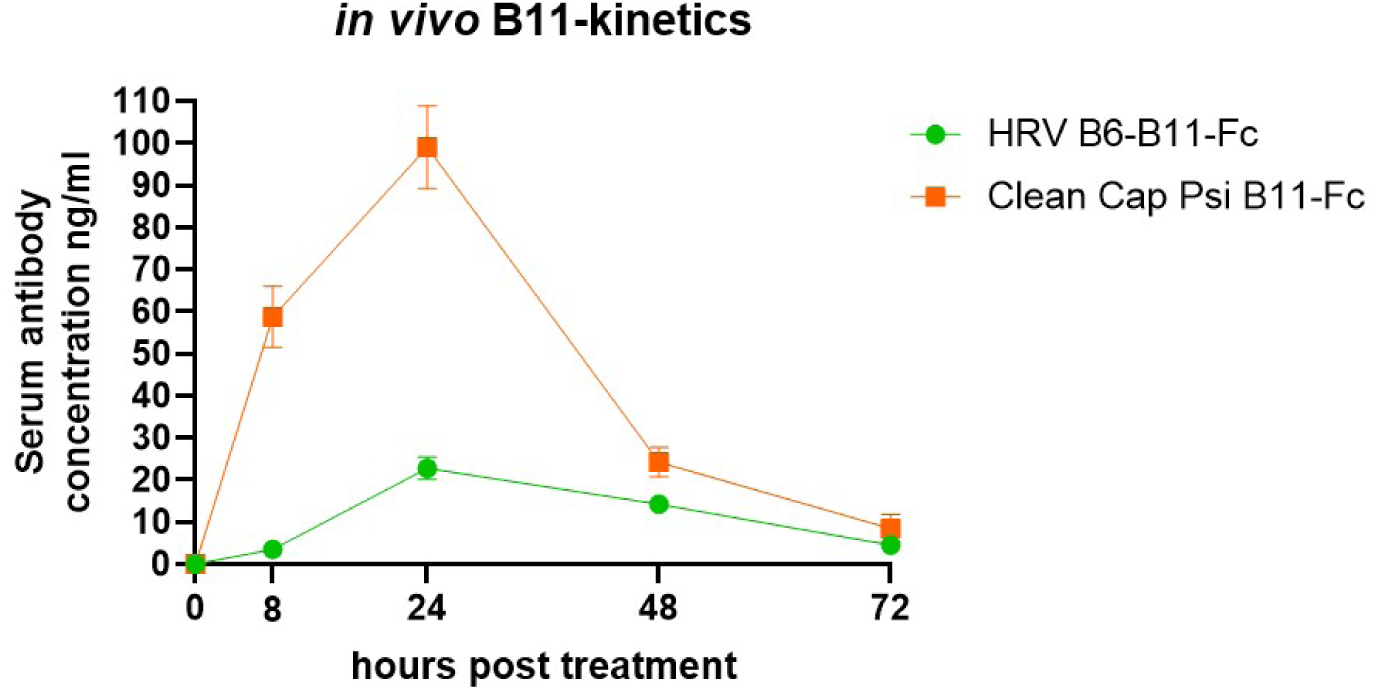
I*n vivo* kinetics curve of B11-Fc antibody expression by the linear and the circular mRNA vectors..

*In vivo* kinetics of B11-Fc expression by mRNA vectors showed a higher level of expression for the linear vectors at all the time points. Maximum values were 99 ±9,8 ng/ml for linear vector and 22,75 ±2,7 ng/ml for the circular vector at the 24-hour time-point.

We then compared the protective activity of the obtained circular vector with the linear one. For this experiment mice were divided into three groups of 16 animals each and were treated with LNPs as in previous experiment. At 4, 16, 24 and 48 hour post-treatment mice were challenged intraperitoneally with 5LD_50_ of BonT/A The survival test results are shown in Figures 8 A – D.

**Figure 8.**
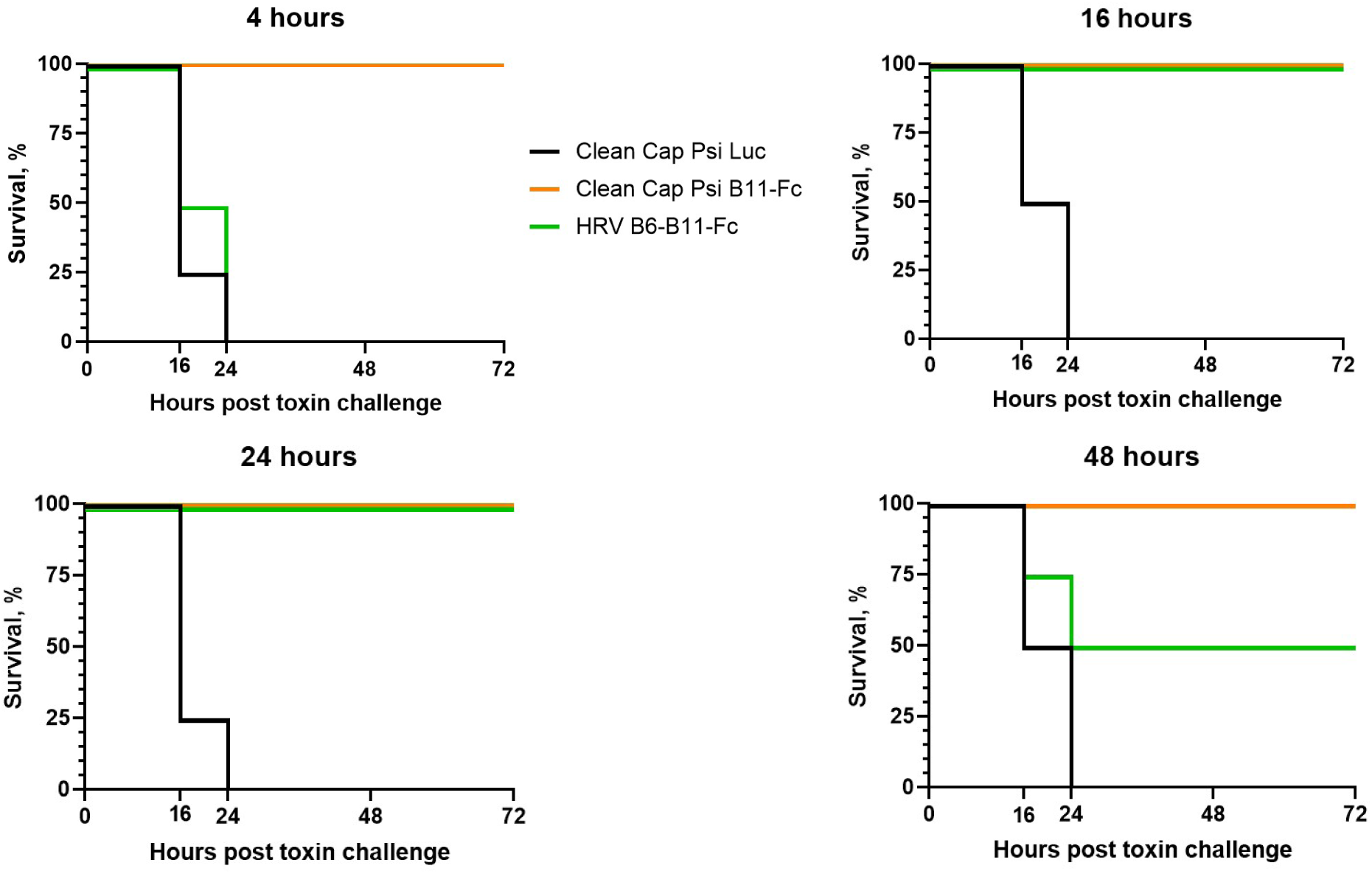
Survival test after 5LD_50_ BonT/A challenge of treated mice. A – mRNA treatment 4 hours before toxin challenge, B – mRNA treatment 16 hours before toxin challenge, C – mRNA treatment 24 hours before toxin challenge, D – mRNA treatment 48 hours before toxin challenge.

All control mice died within 24 hours after toxin injection, while the treated mice showed variable survival. Among the mRNA groups the linear vector showed 100% protection at all time points. The circular vector demonstrated 100% protection at 16- and 24-hour time points, but only 50% protection at 48 hour timepoint. No protection was observed at the 4-hour time point, due to low expression levels.

## Discussion

### *In vitro* luciferase activity

Linear and circular vectors exhibit different expression patterns. Linear vectors reached their maximum at the 16^th^ hour post-transfection, followed by a plateau lasting for about 5 hours and a subsequent dramatic decrease. Circular vectors reached their maximum at the 20^th^ hour post-transfection followed by a plateau lasting nearly 15 hours and then a gradual decrease in luciferase activity. The luciferase assay revealed that linear mRNA containing canonical nucleotides produced low luciferase expression, whereas linear vectors, containing N1-methylpseudouridine demonstrated markedly higher levels of luciferase activity. During the first 24 hours the linear vectors outperformed the circular ones; however, by 48 hours post-transfection the circular vectors surpassed the circular vectors containing N1-methylpseudouridine. Among the linear vectors mRNA constructs capped with CleanCap generated the higher luciferase activity compared to ARCA-capped mRNA.

Finally, the circular mRNA bearing the HRV-B6 IRES exhibited approximately two-folds higher luciferase activity in comparison with the circular vector with CVB3 IRES.

Further kinetics studies were limited by a number of factors. Firstly, CLARIOstar plate reader (BMG Labtech) lacked the required humidity causing the outer wells of the plate to dry slow. Secondly, after five days without reseeding, the cells entered the stationary phase and began detaching from the bottom of the wells. Nevertheless, we anticipate that the decline in luciferase activity of the circular mRNA vectors will be less dramatic than that of the linear vectors over the next few days.

### *In vivo* luciferase activity

As observed, linear mRNA vectors show a more significant increase in luciferase activity in the first hours aftertreatment, which is consistent with the results of the prior experiment. Linear vectors surpass the circular ones in the first 24 hours post-mRNA injection’ By the 48^th^ hour, however, the situation changes dramatically, with circular vectors achieving a higher level of luciferase activity than linear vectors. Surprisingly, the decrease in luciferase activity of the linear vectors is more pronounced *in vivo* compared to *in vitro* study. In contrast, the expression pattern of circular vectors remains comparable to the ones observed in the *in vitro* experiment.

In summary, our luciferase experiments demonstrate that circular vectors exhibit a longer lasting expression in comparison with linear vectors, although their peak expression levels are lower.

### Expression of S-glycoprotein of SARS-Cov2

In accordance with luciferase assay linear vectors with canonical nucleotides showed low, levels of protein expression (114 ng/mL for ARCA-U-S and 153 ng/mL for CleanCap-U-S).

The linear vector, modified with N1-methylpseudouridine and capped with CleanCap remained the highest performer as before. However, the linear vector capped with ARCA was displaced from second place by the circular vector with HRVB6 IRES (519 ng/mL and 875 ng/mL, respectively). These data correlate with findings by Qu L.et.al., who also demonstrated the superiority of circular mRNA over linear mRNA capped with cap0. These data correlate with Qu L. et.al. who also demonstrated superiority of circular mRNA over linear mRNA, capped with cap0 [16].

Surprisingly, expression of S-glycoprotein of SARS-Cov2 by circular mRNA-vector, containing IRES HRVB6 was close to linear CleanCap-Ψ-S (875 ng/mL and 1100 ng/mL, respectively). For linear mRNA modified with N1-methylpseudouridine (m1Ψ the difference between CleanCap-S and ARCA-S was approximately 2 times (1100 ng/mL against 519 ng/mL). The difference between the circular mRNA-vector containing CVB33 IRES and the linear CleanCap-Ψ-S decreased from 16-fold in the luciferase assay to 4.5 fold in antigen expression.

Similarly, the disparity between linear and circular vectors was reduced compared to the luciferase assay. These differences between luciferase and ELISA results may be attributed to a higher stability of the SARS-Cov2 S-glycoprotein in comparison to luciferase, which is an enzyme.

### Immunogenicity study

The results of the immunogenicity study are closer to the expression of the target protein, rather than the luciferase assay. As demonstrated, the immunogenicity of circular mRNA with the IRES of CVB3 is the lowest among the tested vectors. As expected, CleanCap provides better immunogenicity in comparison with ARCA. At the same time, HRV B6 showed immunogenicity, equal to that of the linear mRNA with CleanCap and significantly surpasses mRNA-vector with ARCA. We hypnotised that the difference between the linear vector with CleanCap and the circular vector with IRES HRVB6 would become apparent at lower doses.

### Immunogenicity and protective activity of two “best” vectors

Interestingly, mice, immunised with either linear or circular vectors at a dose of 3 µg per mouse exhibit similar levels of GMT. This finding demonstrates that the linear circular vector with IRES HRVB6 is not only superior to the linear mRNA capped with ARCA reagent but is also equivalent to the widely used mRNA capped with CleanCap and modified with N-1-Me-Pseudouridine.

Our experiment on protective activity also showed that the circular mRNA vector protects mice from lethal infection and significantly reduces viral load. According to RT-PCR, both vaccinated groups had more than 3Log _10_ reduction in viral load in lungs compared to the negative control, with no significant difference between the vaccinated groups.

Viral load measured by TCID_50_ titration on Vero cells yielded similar results; the average TCID_50_ was below the limit of detection for both groups, although the circular vector showed a slightly smaller decrease in viral load.

Our data is consistent with the work of Qu L. et.al. They showed that immunization at a dose of 2.5 µg/mouse provides equal endpoint antibody titers for linear and circular mRNA vectors. In contrast to our study, they performed a virus neutralization test instead of SARS-CoV2 challenge in mice, so it is not possible to completely compare protective activity. Nevertheless, both studies demonstrate in different ways that circular mRNA vaccine is an effective way to protect mice from SARS-CoV2. The distinction lies in the antibody titers for linear and circular mRNA in our work and theirs relates to the differences in the antigen used: we employed the gene of the hole Spike protein instead of RBD domain [16].

It is important to mention the size and structure of our poly-(A) tail. Due to the common problem of poly-(A) deletion in linear mRNA-templates during cultivation we designed an “island” containing 10 other nucleotides, resulting in a poly-(A) tail with a 30/70 ratio as opposed to 100-A poly-(A) used, by Qu L. et.al. which can be deleted during cultivation causing lower results for the linear vector [16].

### Application of circular mRNA-vector for passive immunization

*In vivo* kinetics of antibody expression showed the expected dramatic superiority of the linear vector over the circular vector in the first 24 hours post-treatment. By the 48^th^ hour post-treatment, however, serum antibody concentrations were similar between the two groups.

Unfortunately, the antibody titre produced by the circular vector was near the detection limit at some timepoints, while the linear vector produced a more complete curve. We assume that the linear vector, consistent with luciferase kinetics, produces significant amounts of antibodies in the first hours, followed by a sequential decrease, whereas, the circular mRNA-vector continuously produces lower quantities of antibodies. In our next experiment mice were challenge with 5LD_50_ of BoNT/A. The circular mRNA vector did not reach protective antibody concentrations within the first four hours post-treatment. At 16 hours, however, all mice had reached protective antibody concentrations. This effect was observed at the 24-hour time point. At 48-hour point, the survival in circular mRNA group was 50%, while the linear group had 100% survival. This data correlates with the luciferase assay, where 48-hour activity was approximately 75% of the 24-hour luciferase activity, corresponding to 75% protection from 5LD_50_.

In the course of our research, we demonstrated that our circular mRNA vector is suitable not only for vaccination but also for passive immunization.

## Conclusion

Our work provides a comprehensive comparative analysis of different types of linear and circular mRNA vectors. Linear vectors were produced using either N-1-Me-Pseudouridine or Uridine and two capping reagents: ARCA or CleanCap. Circular vectors contained IRESes elements from Coxackie virus B3 or a novel human Rhinovirus B6.

We demonstrated that the luciferase assay should be considered as a preliminary tool, but should not serve as the sole criterion for comparing vaccine vectors Likewise, expression or immunogenicity studies alone are insufficient. All these assays should be used in combination with protective activity studies for a comprehensive evaluation. Additionally, we obtained a circular vector that provides immunogenicity and protective activity equal to that of the linear vector with CleanCap. Our results provide further evidence that circular mRNA is a relatively new and powerful tool for vaccine development and production, capable of competing with widely used linear mRNAs.

## MATERIALS and METHODS

### Cells and bacterial strains and viruses

*Escherichia coli* DH10B (Invitrogen, USA) was used for DNA preparation. HEK-293 and Vero E6 cells were purchased from ATCC (CRL-1573 and CRL-1586. respectively). Cells were cultured at 37 ^°C,^ in a humidified incubator with 5% atmosphere of CO_2_. Dulbeccòs Modified Eagle Medium (Corning, 10-013-CV) was used for cultivation. HEK-293 medium contained 10% heat inactivated fetal bovine serum (HI-FBS), while Vero cell medium contained 2% HI-FBS.

### mRNA production

DNA-templates for mRNA-production were obtained by Gibson cloning methods using GeneArt™ Gibson Assembly HiFi Master Mix (Invitrogen™, A46628) according to the manufacturer′s instructions. Correct assembly was confirmed via Sanger sequencing. Purified DNA-templates were linearized overnight and complete digestion was confirmed with agarose gel electrophoresis.

mRNA was produced via *in vitro* transcription (IVT) using the HiScribe T7 High Yield mRNA Synthesis Kit (NEB #E2040S) according to the manufacturer′s recommendations. The reaction mix for circular mRNA contained canonical nucleotides and no Cap analogues, while the linear production mix was prepared in different variations: it contained either Uridine-5-Triphosphate or 1-Methylpseudouridine-5-Triphosphate (YEASEN # 10657ES20) and one of two capping reagents the m7G(3’OMe)pppA(2’OMe)pG mRNA cap structure analogue (TriLink®#Т7113) or the anti-reverse cap analogue (Thermo, #AM8045).

The next step for both mRNA-types was treatment with DNase I (New England Biolabs #M0303S) for 30 minutes at 37 °C to digest the DNA templates.

For circular mRNA there was an additional cyclization step. GTP was added to the linear precursor to a final concentration of 2 mM and incubated at 55 °C for 15 minutes to catalyze the cyclization reaction. The mRNA was then purified with the Monarch mRNA Cleanup Kit (New England Biolabs #T2040L). To purify the circular product from residual linear precursor, samples were heated at 65 °C for 3 minutes and cooled on ice before RNase R treatment (Epicenter #RNR07250) at 37 °C for 30 minutes. Final IVT products were purified with the mRNA Clean & Concentrator Kit (ZYMO #R1018).

### In vitro Firefly luciferase assay

HEK-293 cells were seeded into white FB/HB 96-well plates (Greiner) at 2 × 10^4^ per well in 150 µl of DMEM. Six hours later, after cells had finally attached to the bottom, they were transfected with various luciferase-coding mRNAs using with Lipofectamine™ 2000 Transfection Reagent (Invitrogen™) according to the manufacturer’s instructions. The dose of mRNA was 20 ng per well. Fifteen minutes after transfection, D-luciferin (Promega) was added to a final concentration of 5 mM.

Measurement of *in vitro* luciferase kinetics was performed using a CLARIOstar plate reader (BMG Labtech) equipped with an Atmospheric Control Unit. Cells were cultured in 5% CO_2._ at 37 °C. Luciferase activity was measured every hour. All the experiments were performed in triplicate with different IVT-batches and cell passages.

Transfection was performed using Lipofectamine™ 2000 Transfection Reagent (Invitrogen™) as per the manufacturer’s instructions.

### In vitro Expression of SARS-Cov2

HEK-293 cells were seeded to 6-well plates at 3 × 10^5^ cells per well. The next day, when confluency reached 70% cells were transfected with different Spike-coding mRNAs at a dose of 1 µg per well using Lipofectamine™ 2000 Transfection Reagent (Invitrogen™) according to the manufacturer’s instructions. Twenty hours post-transfection cells were collected, washed three times with PBS and lysed. Cell lysates were centrifuged and used for ELISA assay.

The day prior to the analysis, 1p1B10-Fc antibodies were adsorbed onto 96-well plates (Costar, USA), at +4 °C overnight [20]. The quantity of antibodies was 100 ng per well. Then the plate was washed three times with 0.05% Tween-20 PBS (TPBS). After one-hour of blocking and a triple wash with TPBS, cell lysate supernatant and control samples were added. Recombinant S-glycoprotein of SARS-CoV2-Wuhan (Sino Biological, China) was used as a positive control. After one-hour of incubation at 37 °C and five washes with TPBS wash detecting antibodies were added.

As primary antibodies we used 1:1000 diluted polyclonal serum from mice immunized with S-glycoprotein of SARS-CoV2. Detection was performed at 37 °C for one hour followed by five washes with TPBS.

Secondary antibodies were then added with additional one-hour incubation and five TPBS washes. As secondary antibodies we used HPR-conjugated sheep anti-mouse antibodies (NA931V, GE Healthcare, USA).

Finally, TMB substrate (Immunotech, Russia) was added and incubated for 30 minutes at 37°C. The reaction was stopped by addition of 1М H_2_SO_4_. OD_450_ was measured using a CLARIOstar plate reader (BMG Labtech).

### Formulation of mRNA in lipid nanoparticles

Lipid nanoparticles (LNP) assembly procedure was performed as previously described [21]. briefly, lipid nanoparticle included the components, listed below: ionizable lipid (ALC-0315), DSPC, cholesterol and PEG-lipid. Their molar ratio was 46.3:9.4:42.7:1.6. Lipid components were dissolved in ethanol, while purified mRNA was dissolved in 10 mM sodium citrate buffer (pH 3.0) to a final concentration 0.2 mg/mL. Then aqueous and ethanol fractions were mixed in ration 3:1 using a Nanoassemblr Spark device (Precision NanoSystems, USA). Obtained substance was dialyzed against PBS (pH 7.2) with 10% sucrose in Slide-A-Lyzer dialysis cassettes (Thermo Fisher Scientific, USA) for 24 hours. Then formulation was filter sterilized through 0.22 μm Briefly, the lipid nanoparticle formulation included the components, listed below: ionizable lipid (ALC-0315), DSPC, cholesterol and PEG-lipid in a molar ratio of 46.3:9.4:42.7:1.6.

Lipid components were dissolved in ethanol, while purified mRNA was dissolved in 10 mM sodium citrate buffer (pH 3.0) to a final concentration of 0.2 mg/mL. The aqueous and ethanol fractions were mixed in a ratio of 3:1 using a Nanoassemblr Spark device (Precision NanoSystems, USA). Obtained substance was dialyzed against PBS (pH 7.2) with 10% sucrose in Slide-A-Lyzer dialysis cassettes (Thermo Fisher Scientific, USA) for 24 hours.

Then formulation was filter-sterilized through 0.22 μm PES-filter syringe. Prior to use, lipid nanoparticle characteristics were measured: diameter, size distribution and Zeta-potential were evaluated using Zetasizer Nano ZS instrument (Malvern Panalytical) according to the manufacturer′s manual. Total encapsulation efficiency and concentration of mRNA in final formulations were determined by RiboGreen assay (Quant-iT™ RiboGreen™ mRNA Reagent, Thermo Fisher Scientific) as previously described [21].

### SARS-CoV-2 Preparation

The Wuhan-like SARS-CoV-2 virus strain hCoV-19/Russia/Moscow_PMVL-1/2020 was used in the study. SARS-CoV-2 virus was propagated in Vero E6 cells in DMEM with 2% HI-FBS, harvested after 72 hours, aliquoted, titrated on Vero E6 cells and stored at −80 °C. The virus titer was determined on Vero E6 cells using a 50% tissue culture infectious dose (TCID_50_) assay. Serial 10-fold dilutions of the virus stock were prepared in DMEM with 2% HI-FBS and in the volume of 100 μL were added to Vero E6 cells in a 96-well plate in 8 replicates. The cells were incubated at 37 °C in 5% CO_2_ for 96-120 hours and scored visually for cytopathic effect. The TCID_50_ titer was calculated by the Spearman–Kerber method.

### Ethical statement

All animal experiments were approved by the Institutional Animal Care and Use Committee (IACUC) of the Federal Research Centre of Epidemiology and Microbiology named after Honorary Academician N.F. Gamaleya (protocol #93 of May 14, 2025). All procedures with SARS-CoV-2 were carried out in approved biosafety level 3 facilities. All animal experiments were performed in strict accordance with the recommendations of the National Standard of the Russian Federation [22].

### Animal experiments

Protective efficacy was studied in 6–7 week old hemizygous K18-ACE2-transgenic F1 mice obtained by crossing transgenic males B6.Cg-Tg(K18-ACE2)2Prlmn/J, health status SOPF (Jackson Laboratory, USA) and non-transgenic females C57BL/6 Gamrc health status SPF (Gamaleya Research Center, Russia). BALB/c mice were used for *in vivo* luciferase assay and for B11-Fc study. Mice had free access to water and food and were housed in an ISOcage N system (Tecniplast, Buguggiate, Italy).

For the *in vivo* luciferase assay mice were randomly divided into five groups and were treated intramuscularly with LNPs, containing different luciferase-coding mRNAs at a dose of 10 µg per mouse. At 10, 24, 48 and 72 hours post-mRNA treatment, 2.5 µg of D-luciferin was injected intraperitoneally. For SARS-CoV2 studies, mice were immunized twice with S-glycoprotein coding mRNA-vectors at a dose of 3 or once at a dose of 10 µg. As a negative control, luciferase-coding linear mRNA capped with CleanCap reagent and containing N1-methylpseudouridine was used. The interval between immunizations was 21 days. Blood was collected from five mice of each group 21 days after the first of the second immunization using local anesthesia with lidocaine. The size of blood probe did not exceed 100 µl per mouse.

The remaining mice in each group were challenged intranasally at a dose of 10^5^ TCID_50_ with SARS-CoV-2 virus (Wuhan-like). Five animals per group were euthanized on day 4 post-challenge for macroscopic analysis of lung damage and determination of viral load in the lungs. Euthanasia was performed using an overdose of injectable anesthesia. After autopsy the lungs were removed and washed with PBS. A piece of tissue was dissected, placed in homogenization tubes, weighed and supplemented with PBS at 90% of the tissue weight

Another ten animals per group were monitored for weight loss and survival for 21 days. Animals that lost more than 20% of their initial body weight were euthanized before the end of the study.

For the therapeutic potential study of mRNA, mice were treated with 10 µg of B11-Fc-coding mRNA; the negative control remained the same. Then blood was collected at 8, 24, 48 and 72 hours post-injection (three mice pergroup, per time-point)The procedure for blood collection is described above.

Botulinum toxin A was obtained as previously described [23]. The toxin was injected intraperitoneally at a dose of 5LD_50_ at different time-points after mRNA-treatment. Mice were observed for 48 hours. Death time was defined as the time when a mouse was found dead or was euthanized at a humane endpoint by carbon dioxide asphyxiation followed by cervical dislocation.

### In vivo kinetics of B11-Fc expression by linear and circular mRNA vectors

The concentration of recombinant B11-Fc antibody was measured using ELISA kit IgG total-EIA-BEST (Vector-Best, Russia), according to the manufacturer′s instructions, with the following modifications: substitution of kit′s secondary antibodies with ECL Human IgG, HRP-linked whole Ab (Cytiva, NA933-1ML) and use of purified B11-Fc instead of kit′s standard sample.

### Endpoint anti-Spike antibodies GMT measurement

An ELISA protocol was used to evaluate anti-RBD antibodies. Serum samples were purified by centrifugation (800× g, 10 min). Recombinant SARS-CoV-2 RBD domain of B.1.1.1 SARS-CoV-2 virus (SinoBiological, Beijing, China) was used for overnight coating of 96-well plates (100 ng/well). Plates were washed five times with washing solution (PBS + 0.1% Tween-20. TPBS) and then blocked with blocking solution (TPBS with 5% non-fat dry milk). Serum samples were serially diluted in blocking solution and added to wells, followed by incubation at 37 °C for 1 hour. After washing, anti-mouse total IgG secondary HRP-conjugated antibodies (1:5000 dilution, Abcam, Cambridge, UK) in blocking solution were added and plates were incubated at 37 °C for 1 hour. After a final wash, TMB substrate was added and plates were incubated at 20–25 °С. The reaction was stopped with 4 M H_2_SO_4_. The colorimetric signal was measured at 450 nm using a Multiscan FC spectrophotometric plate reader (Thermo Fisher Scientific, Waltham, MA, USA) 30 minutes after the addition of stop solution. The ELISA titer was defined as the highest serum dilution, that produced a colorimetric signal at least twice that of the corresponding dilution of the control serum.

### Determination of Viral Load in Lungs

Ten percent lung homogenates in DMEM with 2% heat-inactivated FBS were prepared on day 4 after challenge using an MPbio FastPrep-24 (MP Biomedicals, Irvine, CA, USA). Homogenates were centrifuged at 12.000× g for 10 minutes and the supernatant were used for further analysis.

The infectious virus titer was determined as described above on Vero E6 cells in a 96-well plate in four replicates after a 120-hour incubation.

mRNA was isolated from virus-containing samples using RNeasy Plus Universal Kits (QIAGEN, Germany) according to the manufacturer’s protocol. RV-RT-PCR was performed using the POLIVIR SARS-CoV-2 “Express” kit (Lytech, Russia) as per manufacturer’s instructions. SARS-CoV-2 viral mRNA with a known concentration of GE/ml was used to construct the calibration curve.

### Statistical analysis and graphical design

All statistical calculations were performed using GraphPad Prism 10. The normality of data distribution was evaluated in the d’Agostino–Pearson test. Comparisons of unpaired samples were performed by the Mann–Whitney test. Multiple data sets were compared by non-parametric ANOVA (Friedman’s test) with Dunn’s multiple comparison post-test. Survival curves were compared using the Log-rank (Mantel–Cox) test.

Visual schemes of the experiments were made using Biorender

## REFERENCES

1. Meo SA, Bukhari IA, Akram J, Meo AS, Klonoff DC. COVID-19 vaccines: comparison of biological, pharmacological characteristics and adverse effects of Pfizer/BioNTech and Moderna Vaccines. Eur Rev Med Pharmacol Sci. 2021;25(3):1663–9. doi: 10.26355/eurrev_202102_24877.

2. Chen H, Liu D, Guo J, et al. Branched chemically modified poly(A) tails enhance the translation capacity of mRNA. Nat Biotechnol. 2024. 10.1038/s41587-024-02174-7

3. Vogel AB, Kanevsky I, Che Y, et al. A prefusion SARS-CoV-2 spike mRNA vaccine is highly immunogenic and prevents lung infection in non-human primates. bioRxiv. Preprint posted online September 8, 2020. doi: 10.1101/2020.09.08.280818.

4. DiPiazza AT, Leist SR, Abiona OM, et al. COVID-19 vaccine mRNA-1273 elicits a protective immune profile in mice that is not associated with vaccine-enhanced disease upon SARS-CoV-2 challenge. Immunity. 2021;54(8):1869–1882.e6. doi: 10.1016/j.immuni.2021.06.018.

5. Liu X, Li Z, Li X, et al. A single-dose circular mRNA vaccine prevents Zika virus infection without enhancing dengue severity in mice. Nat Commun. 2024;15(1):8932. doi: 10.1038/s41467-024-53324-5.

6. Yue X, Zhong C, Cao R, et al. Circ mRNA based multivalent neuraminidase vaccine induces broad protection against influenza viruses in mice. npj Vaccines. 2024;9(1):170. doi: 10.1038/s41541-024-00963-4.

7. Wan J, Wang Z, Wang L, et al. Circular mRNA vaccines with long-term lymph node-targeting delivery stability after lyophilization induce potent and persistent immune responses. mBio. 2024;15(2):e0177523. doi: 10.1128/mbio.01775-23.

8. Zhou J, Ye T, Yang Y, Chuai X, Wang Z, Chiu S. Circular mRNA vaccines against monkeypox virus provide potent protection against vaccinia virus infection in mice. Mol Ther. 2024;32(6):1779–1789. doi: 10.1016/j.ymthe.2024.04.028.

9. Li H, Peng K, Yang K, et al. Circular mRNA cancer vaccines drive immunity in hard-to-treat malignancies. Theranostics. 2022;12(14):6422–6436. doi: 10.7150/thno.77350.

10. Niu D, Wu Y, Lian J. Circular mRNA vaccine in disease prevention and treatment. Signal Transduct Target Ther. 2023;8(1):341. doi: 10.1038/s41392-023-01561-x.

11. Cao J, Novoa EM, Zhang Z, et al. High-throughput 5′ UTR engineering for enhanced protein production in non-viral gene therapies. Nat Commun. 2021;12(1):4138. doi: 10.1038/s41467-021-24436-7.

12. Li T, Liu G, Bu G, Xu Y, He C, Zhao G. Optimizing mRNA translation efficiency through rational 5′UTR and 3′UTR combinatorial design. Gene. 2025;942:149254. doi: 10.1016/j.gene.2025.149254.

13. Asrani KH, Farelli JD, Stahley MR, et al. Optimization of mRNA untranslated regions for improved expression of therapeutic mRNA. RNA Biol. 2018;15(6):756–762. doi: 10.1080/15476286.2018.1450054.

14. Kim S, Sekhon SS, Shin WR, et al. Modifications of mRNA vaccine structural elements for improving mRNA stability and translation efficiency. Mol Cell Toxicol. 2022;18(1):1–8. doi: 10.1007/s13273-021-00171-4.

15. Panova EA, Kleymenov DA, Shcheblyakov DV, Bykonia EN, Mazunina EP, Dzharullaeva AS, et al. Single-domain antibody delivery using an mRNA platform protects against lethal doses of botulinum neurotoxin A. Front Immunol. 2023;14:1098302. doi: 10.3389/fimmu.2023.1098302.

16. Wan J, Wang Z, Wang L, et al. Circular mRNA vaccines with long-term lymph node-targeting delivery stability after lyophilization induce potent and persistent immune responses. mBio. 2024;15(2):e0177523. doi: 10.1128/mbio.01775-23.

17. Qu L, Yi Z, Shen Y, et al. Circular mRNA vaccines against SARS-CoV-2 and emerging variants. Cell. 2022;185(10):1728–1744.e16. doi: 10.1016/j.cell.2022.03.044.

18. McCaffrey AP. mRNA Epitranscriptome: Role of the 5’ Cap. Genet Eng Biotechnol News. 2019;39(5):59–61. doi:10.1089/gen.39.05.17.

19. Chen R, Wang SK, Belk JA, et al. Engineering circular RNA for enhanced protein production. Nat Biotechnol. 2023;41(2):262–272. doi: 10.1038/s41587-022-01393-0.

20. Shcheblyakov DV, Favorskaya IA, Dolzhikova IV, et al. Ultra-potent RBM-specific single-domain antibody broadly neutralizes multiple SARS-CoV-2 variants with picomolar activity. Int J Biol Macromol. 2025;319(Pt 3):145386. doi: 10.1016/j.ijbiomac.2025.145386.

21. Mazunina EP, Gushchin VA, Kleymenov DA, et al. Trivalent mRNA vaccine-candidate against seasonal flu with cross-specific humoral immune response. Front Immunol. 2024;15:1381508. doi: 10.3389/fimmu.2024.1381508.

22. GOST 33044-2014. Principles of Good Laboratory Practice. Interstate Standard. 2014. Available from: https://protect.gost.ru/document.aspx?control=7&baseC=6&page=6&month=2&year=2023&search=&RegNum=1&DocOnPageCount=15&id=238531

23. Derkaev AA, Ryabova EI, Esmagambetov IB, et al. rAAV expressing recombinant neutralizing antibody for the botulinum neurotoxin type A prophylaxis. Front Microbiol. 2022;13:960937. doi: 10.3389/fmicb.2022.960937.

